# Regulatory network and spatial modeling reveal cooperative mechanisms of resistance and immune escape in ER_+_ breast cancer

**DOI:** 10.1101/2025.09.30.679692

**Authors:** Yijia Fan, Sarthak Sahoo, Mohit K. Jolly, Jason T. George

## Abstract

Despite significant progress, the treatment of estrogen receptor-positive (ER+) breast cancer remains clinically challenging due to reversible drug resistance and immune evasion. Drug resistance often arises as cells undergo a dynamic epithelial-to-mesenchymal transition (EMT), while elevated PD-L1 levels contribute to immune escape. While these phenotypic features can variably co-occur, the impact of co-occurrence on the availability of synergistic treatment strategies remains unknown. To investigate their interplay, we constructed an ER-EMT-PD-L1 gene regulatory network and simulated these networks as coupled ordinary differential equations with biologically informed parameters, to generate steady-state expression profiles. Our study revealed that the relevant overarching network generated antagonistic epithelial and mesenchymal modules, capable of producing monostable, bistable, and tristable dynamics. We further examined the link between phenotypes and immune evasion by quantifying average PD-L1 expression, and found that epithelial-sensitive states consistently exhibited low PD-L1. In contrast, hybrid- and mesenchymal-resistant states were associated with high PD-L1, highlighting a strong coupling between EMT, resistance, and immune evasion. Extending on these network-level insights, we further used a spatially explicit agent-based model seeded with GRN-derived phenotypes to probe tumor behavior under therapeutic pressure. Simulations revealed that tumor escape required co-occurrence of therapy resistance, motility, and immune suppression, with plasticity and multistability further promoting adaptive persistence. Lastly, we identified combination therapies predicted to constrain malignant diversification and enhanced immune accessibility. Taken together, our modeling work links regulatory dynamics with tumor-level adaptation and underscores potential strategies to therapeutically reprogram cell states toward sensitivity.

## Introduction

Estrogen receptor-positive (ER+) breast cancer is the most common subtype, accounting for ~80% of cases (1). Although targeted molecular therapies such as tamoxifen, aromatase inhibitors, and fulvestrant have markedly improved outcomes, resistance remains a major clinical problem, especially in advanced disease (2). Progression to late-stage ER+ disease is frequently accompanied by genomic instabil-ity, clonal diversification, and phenotypic plasticity, making tumors biologically distinct and more difficult to treat (3). Mechanisms of resistance in ER+ breast cancer, as in many cancers, can be broadly categorized into genetic alterations and acquired adaptive responses (2, 4). Genetic alterations include constitutively active ESR1 variants and mutations activating pathways such as PI3K/AKT/mTOR (5). Acquired adaptations, by contrast, often involve phenotypic programs such as the epithelial-to-mesenchymal transition (EMT), which confers cellular plasticity and drug tolerance, and may be accompanied by immune evasion strategies such as PD-L1 up-regulation (2, 6). While such variety in adaptive behavior has been observed empirically, the extent to which these phenotypic features co-occur, and the extent to which their co-occurrence shapes tumor behavior and therapeutic efficacy, remains incompletely understood.

Here, we constructed an integrated ER-EMT-PD-L1 Gene Regulatory Network (GRN) and simulated its dynamics using coupled ordinary differential equations with parameters sampled from biologically relevant ranges. Our framework revealed antagonistic epithelial and mesenchymal modules capable of generating mono-, bi-, and tri-stable dynamics, uncovering a strong coupling among EMT, endocrine treatment resistance (specifically resistance to tamoxifen, a selective estrogen receptor modulator widely used as standard endocrine therapy in ER+ breast cancer), and elevated PD-L1 expression (7, 8). To evaluate the clinical relevance of these predictions, we analyzed TCGA ER+ breast cancer cohorts using EMT, tamoxifen-resistance, and PD-L1-associated signatures, and observed consistent positive correlations. Building on our observations that EMT, resistance, and immune evasion are interlinked drivers of tumor adaptation, we further hypothesized that reversing EMT through mesenchymal-to-epithelial transition (MET) might represent a rational therapeutic strategy. Patient stratification using the transcriptomic signature of all-trans retinoic acid (ATRA), a MET inducer, further revealed that mesenchymal-like groups experienced significantly poorer survival. Finally, we embedded GRN-derived phenotypes into a spatially explicit agent-based model (ABM) to examine tumor behaviors under combinational therapy. These simulations demonstrate that escape required the convergence of chemoresistance, motility, and immune suppression, reinforced by plasticity and multista-bility, and further identified combination strategies predicted to constrain malignant diversification and enhance immune accessibility. Taken together, our study integrates regulatory modeling, patient transcriptomics, survival data, and spatial dynamics to link resistance mechanisms with tumor adaptation, offering a computational framework for identifying opportunities to reprogram resistant tumors toward therapeutic sensitivity.

## Results

### Multistability of the ER-EMT-PD-L1 regulatory network drives phenotypic heterogeneity and promotes biased transitions toward targeted therapy resistance and immune-evasive states

To investigate the interplay between resistance to targeted molecular therapy and immune evasion in ER+ breast cancer, we focused on the connections linking core EMT regulators with the ER alpha (ER*α*) isoforms and the immune checkpoint protein PD-L1. The emergence of reversible resistance to therapies like tamoxifen in ER+ breast cancer can be driven by key EMT players including ZEB1, miR-200, and SLUG, which modulate ER*α* activity (9). A switch from the epithelial (E) state towards either the hybrid E/M or mesenchymal (M) state often correlates with acquired tamoxifen resistance and reduced ER*α* levels. Furthermore, given that ER*α* can directly repress PD-L1 transcription, anti-estrogen therapy may inadvertently promote immune evasion by increasing PD-L1 (10).

To explore the emergent dynamics of this system, we constructed a GRN incorporating these key regulatory elements (Fig 1A). The network includes ER*α*66, the primary target of ER blockade therapy, and ER*α*36, a known mediator of resistance (11, 12). Interactions between epithelial-mesenchymal regulators and the ER*α* isoforms were included as previously modeled (9). Furthermore, the gene product of LOXL2 has been implicated in EMT which has important implications for matrix remodeling during EMT(13, 14). We included the following links based on available literature evidence to integrate LOXL2 as an EMT player in the gene regulatory network. LOXL2 is activated by ZEB1 (15, 16), and is suppressed by miR-200 (15). LOXL2 is also known to promote EMT by activating SNAI2 (SLUG) (17, 18), a key partial inducer of EMT (19), as has been previously reported. To generate *in-silico* steady-state gene expression values for our minimal working GRN, we apply RACIPE (20), which simulates a given gene regulatory network as a set of coupled ordinary differential equations, with parameters sampled from biologically relevant ranges. The ensemble of RACIPE-derived steady states is indicative of the possible phenotypes allowed by the network topology. To quantify the cellular phenotype of a given steady-state solution, we defined an EM score from (z-normalized) expression values of ZEB1, SLUG, miR-200, and CDH1. In this context, larger EM scores correspond to enrichment of the mesenchymal phenotype. We similarly defined a resistance score from the expression of the different isoforms of the ESR1 gene (see Methods).

**Fig. 1.**
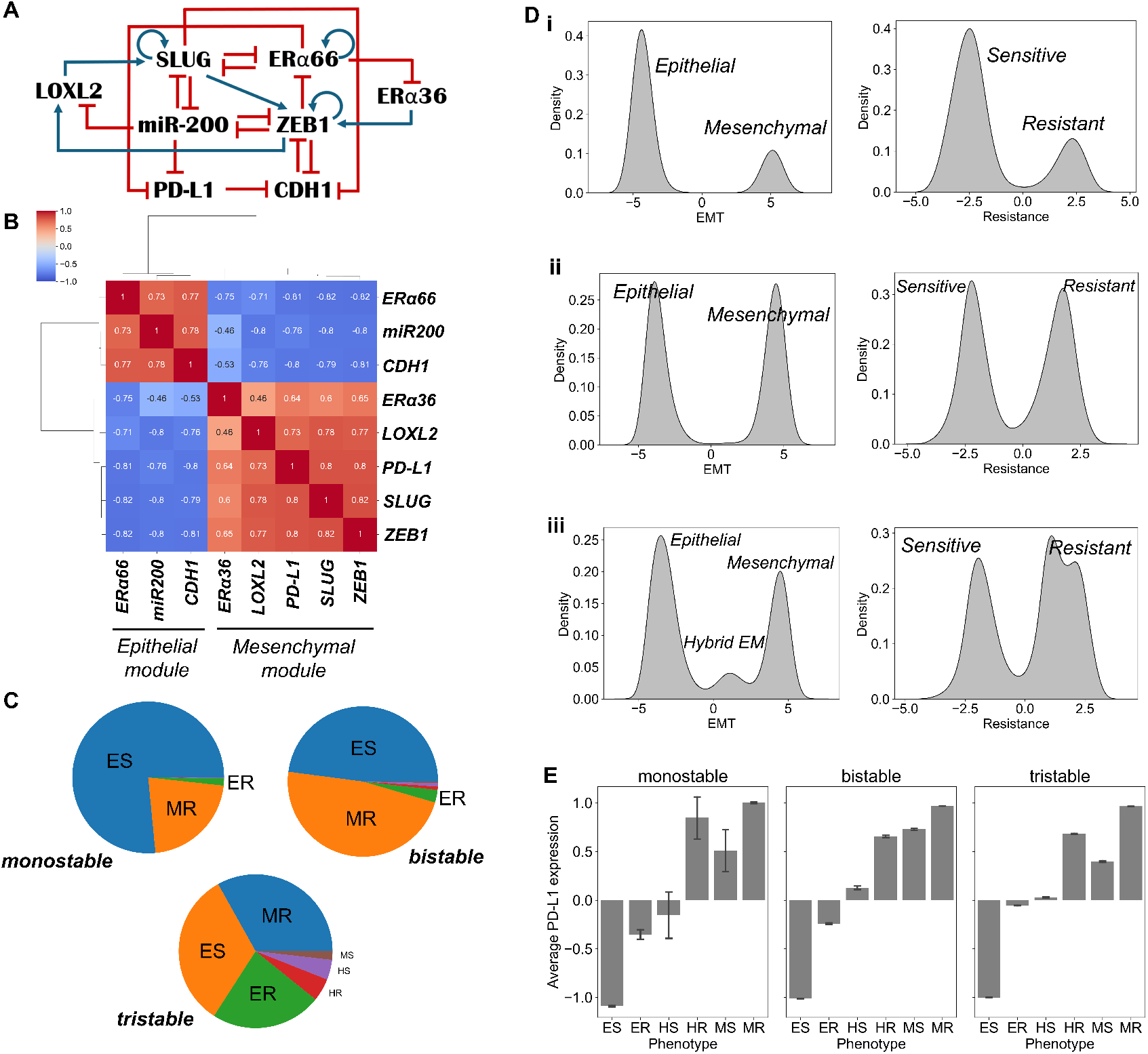
Phenotypic heterogeneity arising from the EMT-ER*α*-PD-L1 regulatory network. (A) Schematic of the GRN coupling core EMT regulators, Estrogen receptors with PD-L1. Blue arrows indicate activation, and red hammer-headed lines represent inhibition. (B) Simulated gene-gene spearman correlation clustermap showing epithelial and mesenchymal modules. (C) Pie charts showing the proportion of cellular phenotypes obtained from parameter sets that result in monostable, bistable, and tristable systems. (D) Kernel Density Estimate (KDE) plots of phenotypes characterized by EMT and Resistance scores for (i) monostable, (ii) bistable, and (iii) tristable parameter sets. (E) Average PD-L1 expression levels for each corresponding phenotype across monostable, bistable, and tristable parameter cohorts. Error bars represent the standard deviation of gene expression.

Simulating this network revealed predominantly two distinct, mutually antagonistic modules. One ‘mesenchymal’ module comprised the EMT-inducers ZEB1, SLUG, LOXL2, along with ER*α*36 and PD-L1 (Fig 1B). These factors were all highly correlated with one another and negatively correlated with the members of the opposing ‘epithelial’ module, which consisted of CDH1, miR-200, and ER*α*66 (Fig 1B). Furthermore, steady-state solutions generated from a large ensemble of randomized parameter sets revealed the capacity of this network to produce monostable, bistable, and tristable dynamics. For parameter sets resulting in a single stable state (monostability), the system was dominated by an Epithelial-Sensitive phenotype, with a smaller but significant fraction of cells adopting a Mesenchymal-Resistant (MR) state (Fig. 1C). Parameter sets that permitted two stable states (bistability) predominantly gave rise to coexistence between the Epithelial-Sensitive (ES) and MR phenotypes (Fig. 1C). The emergence of three stable states (tristability) allowed for a much richer phenotypic landscape, with sizable populations of not only ES and MR cells, but also Epithelial-Resistant (ER), hybrid E/M-Sensitive (HS), and hybrid E/M-Resistant (HR) phenotypes (Fig. 1C). These distinct phenotypes were defined based on their corresponding EMT and Resistance scores, which showed characteristic distributions for each stability regime (Fig. 1D).

We then investigated the inter-dependencies between the coupled (EMT/drug-susceptibility) phenotypes and immune evasion by quantifying the average PD-L1 expression for each state (Fig. 1E). A clear trend emerged across all stability regimes. The ES phenotype consistently displayed the lowest levels of PD-L1. In stark contrast, the HR and MR phenotypes exhibited the highest PD-L1 expression, indicating a strong coupling between the acquisition of resistance, a mesenchymal-like state, and an immune-evasive phenotype. Interestingly, the ER phenotype showed intermediate levels of PD-L1, higher than the ES cells but notably lower than the fully MR cells. This suggests a graded increase in immune evasion potential as cells transition from a Sensitive to a Resistant state, which can occur even without a complete EMT.

### Integration of EMT, tamoxifen resistance, and PD-L1 signatures reveals co-occurrence and prognostic impact

Building on our GRN predictions, we next analyzed transcriptomic signatures from The Cancer Genome Atlas (TCGA) ER+ breast cancer cohort to evaluate whether the predicted couplings among EMT, tamoxifen resistance (TamRes), and PD-L1 are reflected in patient tumors. Single-sample gene set enrichment analysis (ssGSEA) was performed using hallmark EMT gene signatures from the Molecular Signatures Database (MSigDB) (21), a TamRes signature derived from proteomic analysis of resistant MCF7 cells (22), and a publicly available PD-L1 signature (10). Patients were stratified according to their signature scores, with EMT scores partitioned into upper (M), middle (H), and lower (E) tertiles, and TamRes and PD-L1 scores classified at their respective median values (see Methods). Stratification of patients revealed clear phenotypic trends consistent with the *in silico* findings (Fig. 2A-C). We further performed pair-wise correlation analysis across all ER+ samples and found strong positive associations among EMT, TamRes, and PD-L1 signatures (Fig. 2E), supporting their coordinated activation. In addition, we examined the distribution of PD-L1 scores across composite phenotypic groups defined by combined EMT and TamRes states (Fig. 2F), which showed that PD-L1 expression was consistently highest in hybrid- and mesenchymal-resistant groups, reinforcing the link between immune evasion and therapy-resistant phenotypes. Given that EMT, TamRes, and immune evasion appear to act in context as interlinked drivers of tumor adaptation, we further posited that promoting a MET to reverse EMT could represent a rational therapeutic strategy. To assess this, we derived an ATRA-response score based on gene expression changes observed in MCF7 (responsive) and BT474 (resistant) cells before and after retinoic acid (RA) treatment, calculated as a z-normalized difference between responsive and resistant gene sets (see Methods) (Fig. 2D) (23). Importantly, stratification of patients by this ATRA-response signature revealed that resistant-like groups experienced markedly poorer survival (Fig. 2G), consistent with the hypothesis that loss of MET-inducing capacity exacerbates malignant progression. Together, these data validate the GRN-derived predictions, highlight the prognostic value of targeting EMT- and PD-L1-associated programs, and motivate further investigation into how these cooperative phenotypic traits influence tumor-level behavior under therapeutic pressure.

**Fig. 2.**
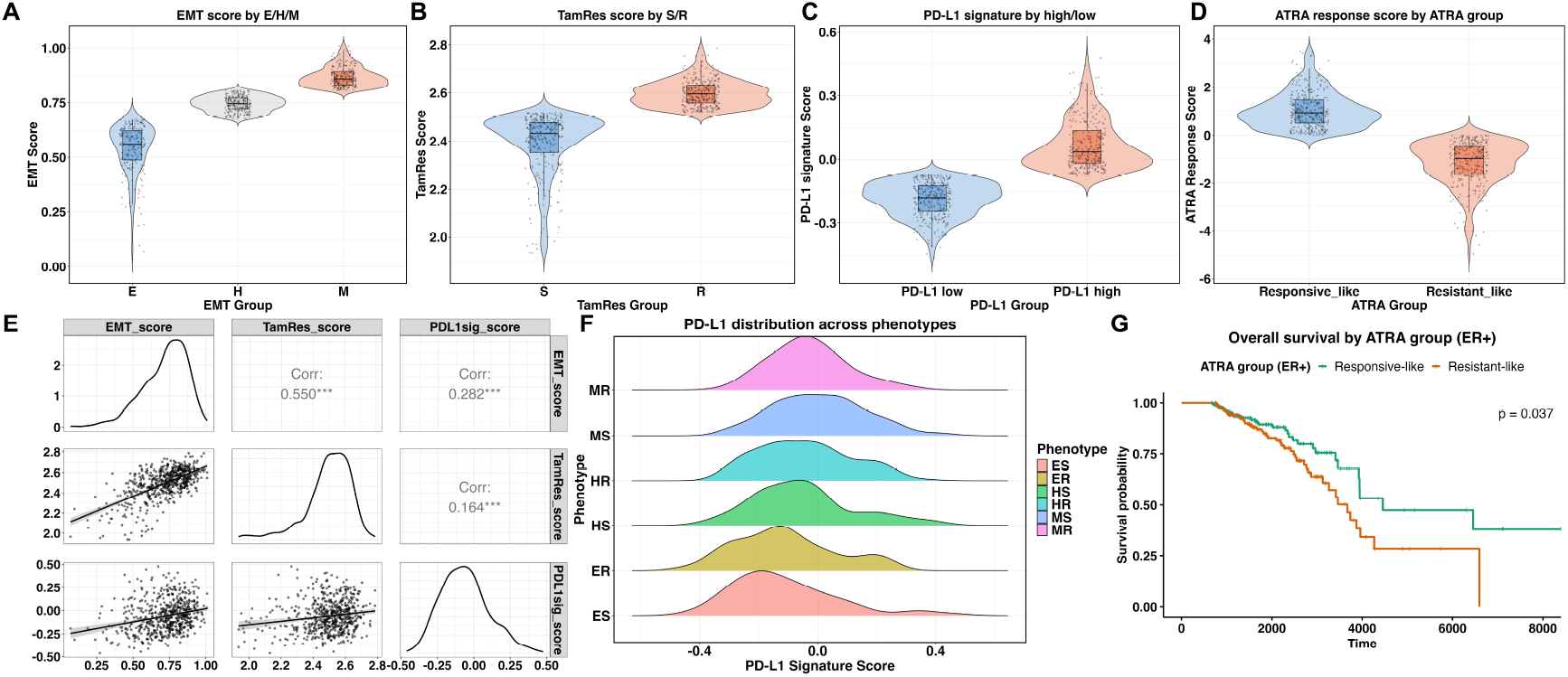
Strong coupling among EMT, TamRes, and PD-L1 signatures in TCGA ER+ breast cancer cohorts, with poorer survival in the ATRA-resistant group. (A) Distribution of EMT scores across E, H, M phenotypes. (B) TamRes scores stratified by S and R groups. (C) PD-L1 signature scores comparing PD-L1 low versus PD-L1 high groups. (D) ATRA response scores stratified by ATRA groups (Responsive-like vs Resistant-like). (E) Pairwise correlation analysis among EMT, TamRes, and PD-L1 signature scores, with Spearman correlation coefficients shown. Asterisks indicate statistical significance of the correlation (*** p < 0.001, ** p < 0.01, * p < 0.1) (F) Distribution of PD-L1 signature scores across composite phenotypes. (G) Kaplan–Meier survival analysis comparing the top 40% of patients in each group, showing poorer outcomes in the Resistant-like group.

### Non-plastic tumor subpopulations utilize functionally distinct but coordinated escape strategies driven by regulatory network architecture

To determine how each phenotype independently contributes to treatment resistance, we investigated the spatial and temporal behaviors of GRN-derived cell states under therapeutic pressure. To this end, we specialized and applied a previously developed spatially explicit ABM (24), seeding tumor cells with a small subset of fixed phenotypes randomly sampled from the RACIPE simulation output. In this model, individual tumor cells and cytotoxic T cells (CD8^+^ T cells), hereafter referred to as T cells, are represented as discrete agents, each with pres-pecified behavior. Tumor cells undergo random migration and proliferation, while T cells migrate chemotactically toward tumor-derived chemokines, kill antigen-expressing tumor cells within an interaction radius, and expand clonally (24–26) (see Methods). We began initially by modeling tumors with static phenotypes lacking plasticity to isolate the role of static heterogeneity in shaping the treatment response. We assumed a default treatment consisting of targeted molecular therapy using tamoxifen (8), and adaptive cytotoxic T cell killing, which together represent the standard combination of endocrine therapy and immune surveillance in ER+ breast cancer. In taking this approach initially, we were able to distinguish the independent contributions of each set of coupled phenotypes to model outcomes, which were then studied later under dynamic state transitions.

Toward this end, we designed three tumor configurations by sequentially introducing key traits derived from GRN analysis. Each tumor cell in the ABM inherits a set of phenotype-associated scores extracted from GRN steady states, including a resistance score (RS), an epithelial-mesenchymal score (EM) score, and a PD-L1 score. In the ABM, RS correlates positively with tumor survival under tamoxifen treatment; EM score indicates the degree of EMT, which relates to increased motility and reduced antigenicity; PD-L1 score reflects the immune evasion capacity against T cell mediated killing (see Methods). We first simulated tumors where cells were characterized only by RS, then next added EMT features on top of RS (RS + EMT), and then finally included PD-L1 expression (RS + EMT + PD-L1). To quantify tumor and immune dynamics, we recorded tumor burden and T cell counts over time (Fig 3A). To further examine phenotypic characteristics and tumor fitness, we also tracked the temporal evolution of resistance score (RS), cell motility (measured by mean displacement), spatial distribution across tumor configurations (Fig 3D-F).Tumors composed solely of RS scores cells remained spatially compact with limited immune evasion (Fig 3A, D). Although resistant to tamoxifen, these cells were vulnerable to immune surveillance and were eventually eliminated by T cells. Introducing EMT features increased overall resistance, motility, and spatial dispersion, producing more diffuse tumor patterns that impaired T cell access and prolonged tumor persistence (Fig 3A-C, E). The addition of PD-L1 further enhanced immune evasion by directly inhibiting T cell cytotoxicity. These tumors exhibited widespread invasion into surrounding regions, with T cell accumulating predominantly at the periphery, indicating spatial exclusion (Fig 3A-C, F). Assessing the resulting correlations between each phenotype of scoring cells revealed strong positive associations among all three phenotypes (Fig 3G). EMT showed a robust correlation with both resistance and PD-L1 expression, suggesting that EMT may co-occur with increased immune evasion and therapy resistance, and that tumors with elevated immune checkpoint expression tend to exhibit greater resistance to therapy, possibly due to an immunosuppresive microenvironment. In addition to exhibiting consistency with the GRN predictions, these findings are also consistent with previous studies (27). Together, these results highlight that effective tumor escape occurs in the setting of the co-occurrence of multiple phenotypic traits, including targeted therapy resistance, high motility, and immune suppression, even in the absence of phenotypic plasticity. While therapy resistance offers protection against drug-induced cytotoxicity, EMT enables cells to dynamically reposition and evade immune predation, high PD-L1 expression provides protection against internal immune surveillance. These traits frequently act synergistically to reinforce tumor persistence.

**Fig. 3.**
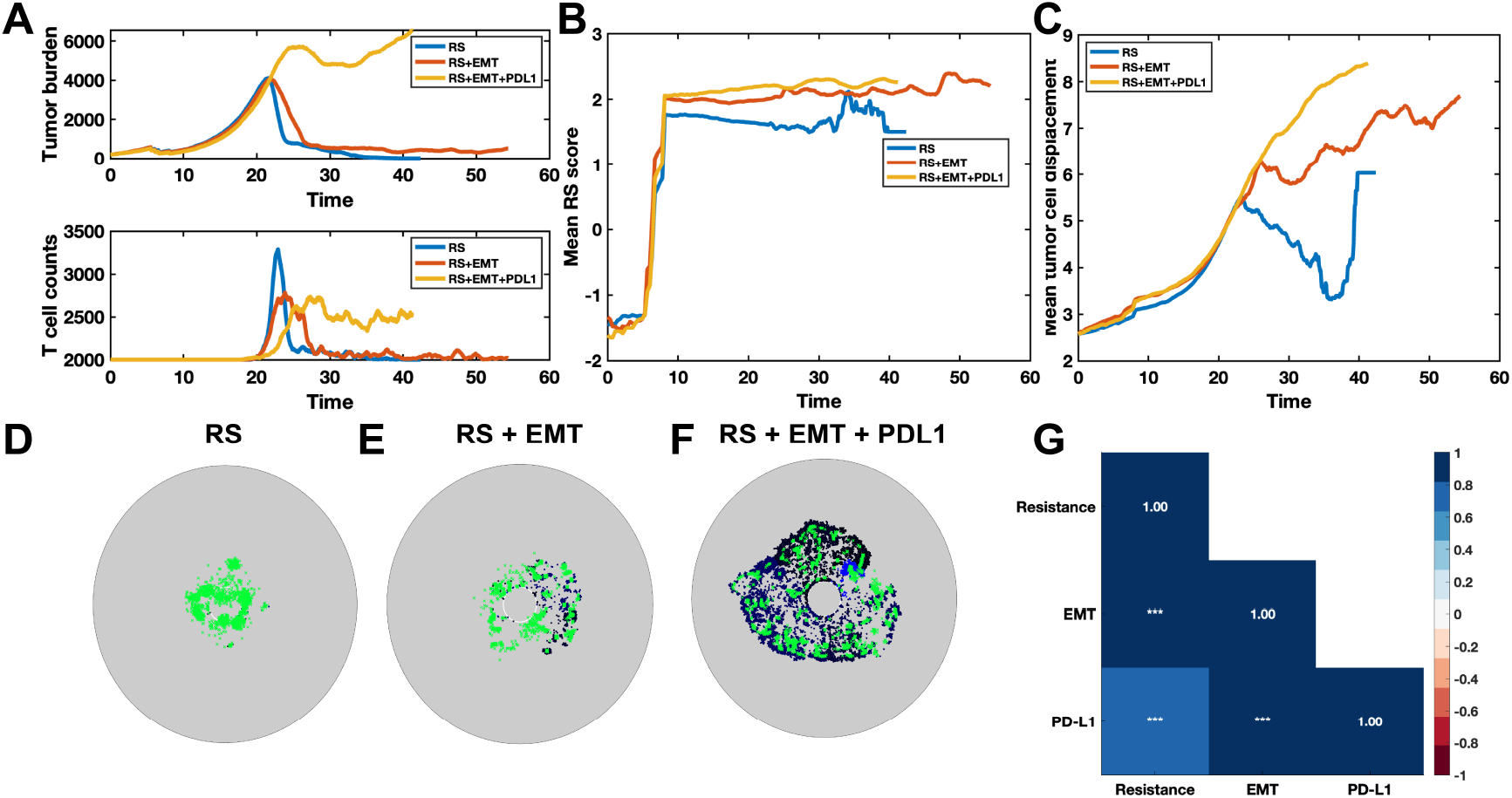
Distinct tumor escape strategies emerge from fixed phenotypic traits under tamoxifen and adaptive immune killing. (A) Time evolution of tumor burden (top) and total T cell count (bottom) under each phenotypic configuration. Tumors with only RS cells were initially reduced but later controlled by T cells, while RS + EMT and RS + EMT + PDL1 tumors exhibited persistent survival and immune suppression. (B) Mean resistance score of tumor cells over time. All groups show rapid selection for high resistance phenotypes, with RS + EMT + PDL1 tumors maintaining the highest overall resistance. (C) Mean displacement of tumor cells over time. EMT increases motility, and the addition of PDL1 further promotes outward expansion. (D-F) Spatial distribution of tumor cells (dark blue) and T cells (green) at a matched time point for the RS (D), RS + EMT (E), and RS + EMT + PDL1 (F) groups. Tumor cells are color-coded in shades of blue according to their epithelial-mesenchymal (EM) score: darker blue indicates stronger mesenchymal characteristics, while lighter blue indicates more epithelial-like phenotypes. RS tumors remain compact and immunologically accessible. RS + EMT tumors exhibit greater spread, and RS + EMT + PDL1 tumors form a large, invasive structure with T cells excluded to the periphery. (G) Correlation matrix of GRN-derived phenotypic scores (Resistance, EMT, and PD-L1) across all stable tumor states. Color intensity reflects Pearson correlation coefficients, with positive correlations shown in blue and negative correlations in red. Asterisks indicate statistical significance (***p < 0.001).

### Tumor plasticity increases adaptability by supporting flexible transitions and survival under combination therapy pressure

To further investigate how phenotypic plasticity interacts with the underlying regulatory structure to shape treatment outcomes, we next extended our simulations to compare tumors governed by monostable, bistable, and tristable GRNs. These regimes differ in the number of coexisting stable gene expression states that are supported by the network: monostable networks permit only one dominant phenotype, bistable networks allow switching between two distinct phenotypes, and tristable networks provide access to three phenotypic states. While previous sections focused on fixed combinations of resistance, EMT, and PD-L1 expression, here we explored how the size and structure of the accessible phenotypic landscape influences adaptive tumor behavior under default combinational treatment (targeted molecular therapy tamoxifen and adaptive T cell killing). In all cases, we initialized the system with a small, randomly selected subset of phenotypes from RACIPE output and allowed dynamic transitions according to GRN-defined switching rates (see Methods). This design enabled us to isolate the impact of multistability on plastic adaptation and treatment response.

Across all conditions, we tracked tumor burden as a primary measure of treatment efficacy (Fig. 4A). Tumors governed by monostable GRNs exhibited the slowest growth and highest treatment sensitivity. In contrast, tumors in the bistable and tristable regime showed markedly accelerated expansion, reflecting enhanced adaptation to dual therapeutic pressure. This result suggests that multistability, when coupled with plasticity, enables tumors to more effectively explore and settle in survival-promoting phenotypes. To further assess these dynamics, we analyzed the temporal evolution of phenotypic composition. We found that tumors with different stability landscapes exhibited distinct rates of transition toward the more aggressive, high-fitness MR state: tristable tumors progressed most rapidly, followed by bistable tumors, while monostable tumors transitioned the slowest (Fig 4B-D). These differences were also evident in the spatial distribution of tumor and immune cells at the treatment endpoint (Fig 4E-G). Monostable tumors remained partially centralized with higher T cell infiltration. Both bistable and tristable tumors exhibited limited immune infiltration, lacking full penetration of T cells into the tumor interior. To better understand the adaptive shifts underlying these patterns, we quantified phenotypic transitions throughout the simulation in both bistable and tristable conditions respectively (Fig 4H-I). In bistable tumors, the most dominant transitions were from ES to MR, indicating a continual drive toward more resistant, mesenchymal phenotypes. In contrast, tristable tumors exhibited ES to ER and ES to MR transitions as the most frequent events.Collectively, these findings indicate that phenotypic plasticity, when combined with regulatory multistability, promotes tumor persistence through dynamic and directional adaptation. While targeted therapy and immune pressure can suppress sensitive cells, high-plasticity systems enable tumors to reconfigure their phenotype distribution in response to stress. Thus, tumor escape under treatment is not merely a result of static heterogeneity, but of the dynamic accessibility and navigability of the phenotypic landscape.

**Fig. 4.**
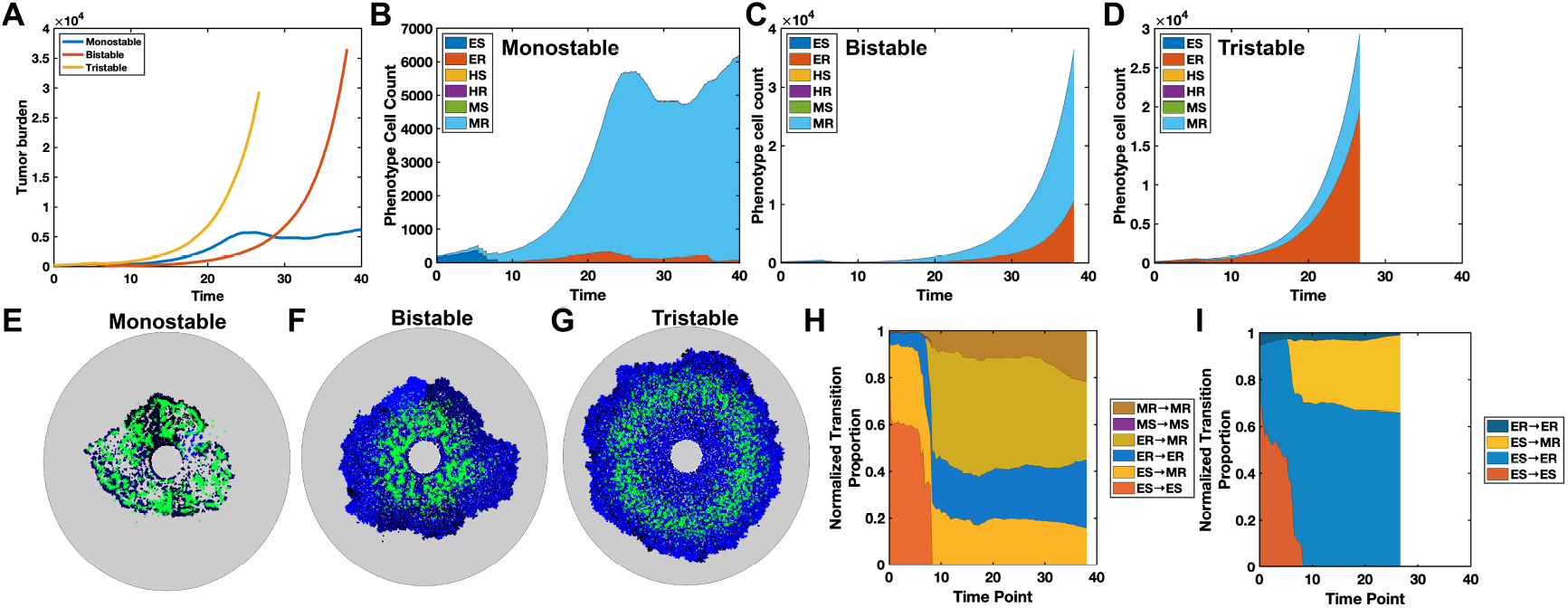
Phenotypic plasticity and GRN multistability together shape tumor evolution under targeted molecular therapy and adaptive immune killing. (A) Tumor burden over time across the three GRN regimes. Tristable tumors exhibited the fastest expansion and treatment evasion, while monostable tumors were most effectively controlled. (B-D) Time evolution of phenotypic composition for monostable (B), bistable (C), and tristable (D) tumors. ES: epithelial-sensitive; ER: epithelial-resistant; HS: hybrid-sensitive; HR: hybrid-resistant; MS: mesenchymal-sensitive; MR: mesenchymal-resistant. Tristable tumors transitioned into mesenchymal phenotypes more rapidly than bistable tumors. Monostable tumors retained a sustainable proportion of HR phenotypes throughout treatment. (E-G) Spatial distribution of tumor (dark blue) and T cells (green) at matched late time point for the monostable (E), bistable (F), and tristable (G) conditions. Tumor cells are color-coded in shades of blue according to their epithelial-mesenchymal (EM) score: darker blue indicates stronger mesenchymal characteristics, while lighter blue indicates more epithelial-like phenotypes. Tristable tumors exhibit widespead expansion and immune exclusion. Monostable tumors remain spatially constrained. (H-I) Normalized transition proportions over time for bistable (H) and tristable (I) conditions.

### Tumor escape dynamics under different treatment combinations are shaped by plastic adaptation and selective transitions

Given that tumor plasticity promotes adaptation under multidimensional pressure, we next tested how combination therapy influences phenotypic transition and treatment outcomes. Specifically, we explored seven treatment regimes combining tamoxifen (denoted as A), ATRA (B), and PD-1/PD-L1 blockade (C), selected to represent clinically relevant strategies targeting hormone signaling, cell differentiation, and immune evasion, respectively (Fig 5A) (7, 28, 29). Treatment-dependent differences in tumor burden dynamics were evident (Fig 5B). While all monotherapies delayed progression to varying degrees, tamoxifen (A) was the most effective among them. However, none were sufficient to control long-term tumor outgrowth. Among dual therapies, tamoxifen combined with ATRA (A+B) produced a synergistic and sustained reduction in tumor burden. Adding PD-1/PD-L1 blockade to tamoxifen (A+C) delayed progression more than PD-1/PD-L1 blockade alone, suggesting that tamoxifen effectively eliminated sensitive tumor cells, but the remaining resistant cells still escaped T cell control. The combination of ATRA and PD-1/PD-L1 blockade (B+C) was more effective than either agent alone, likely due to ATRA-driven reprogramming of malignant, mesenchymal tumor cells into more epithelial, low-PD-L1-expressing phenotypes that were more susceptible to T cell-mediated killing. However, spatial heterogeneity and delayed T cell infiltration allowed continued tumor expansion, resulting in transient states of dynamic equilibrium. The most profound and durable suppression was achieved with the triple combination (A+B+C), reflecting cooperative targeting of hormone signaling, differentiation state, and immune evasion.

**Fig. 5.**
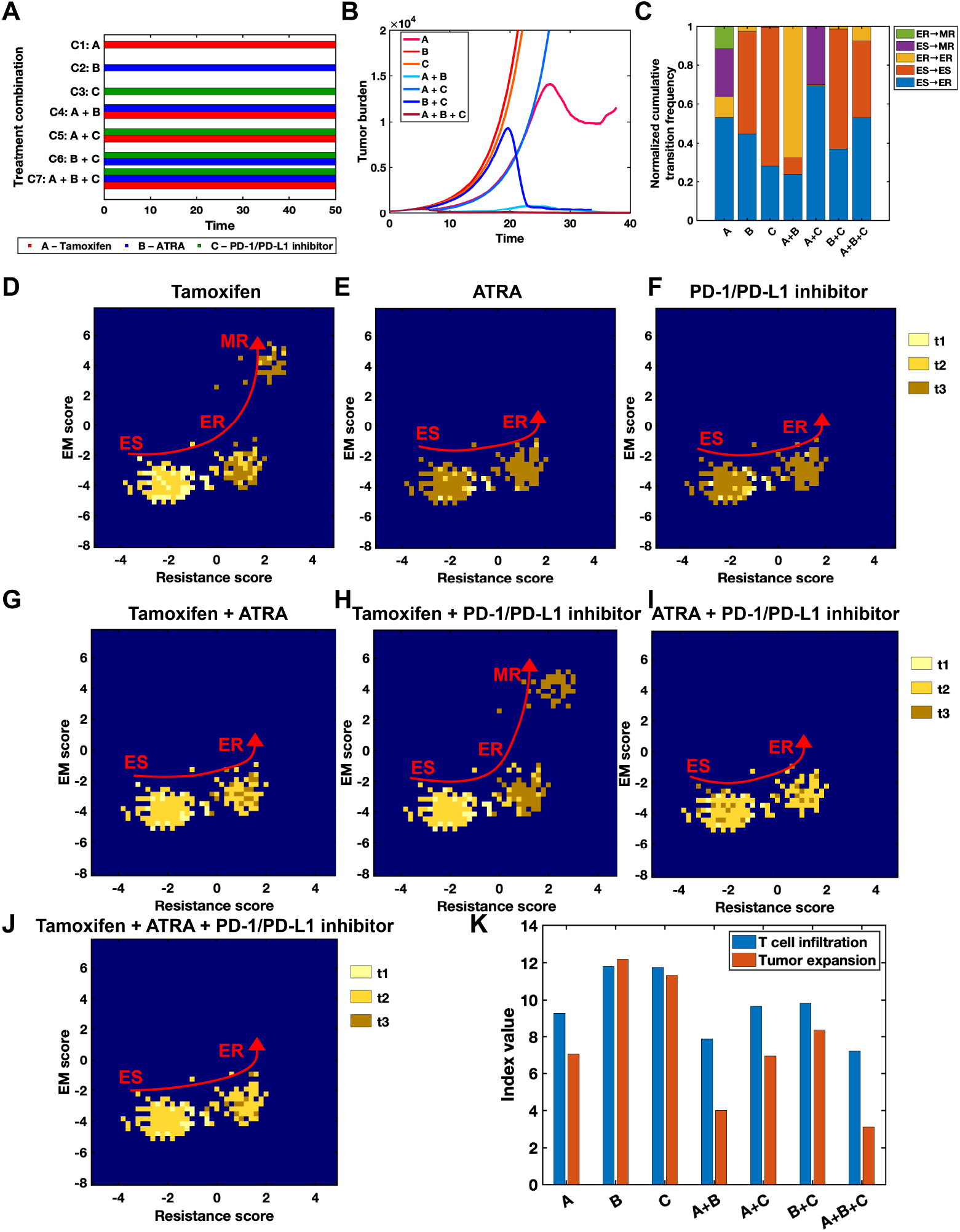
Spatial, phenotypic, and evolutionary dynamics of tumors under different treatment combinations. (A) Treatment timeline of different treatment conditions, each represented by a combination of Tamoxifen (A), ATRA (B), and PD-1/PD-L1 inhibitor (C). (B) Tumor burden over time under differernt treatment conditions. Dual and triple treatment combinations suppress tumor growth to varying degrees. (C) Normalized cumulative phenotype transition frequencies for each treatment conditions. Tumor phenotypic transitions exhibit distinct adaptation patterns, directions, and frequencies under differernt treatment conditions. (D-J) Phenotypic distribution over resistance and EMT score space at three time points (t1, t2, t3) for each treatment condition. Colorbar indicates density over time. Arrows trace dominate phenotypic evolution paths, showing trajectories from epithelial-sensitive (ES) to resistant or more malignant states. (K) Quantification of T cell infiltration and tumor expansion across different treatment combinations. Both the T cell infiltration index (blue) and the tumor expansion index (orange) were calculated as the mean distance of T cells and tumor cells respectively to the center at the end time point of the simulation.

We next quantified cumulative phenotypic transitions under different treatment regimens and examined the distribution of tumor cell states over resistance and EMT score space at three time points (t1, t2, t3) to assess how therapeutic pressure shapes both the direction and spatial organization of tumor evolution (Fig 5C-K). Transition frequencies were normalized across time and treatment groups, revealing distinct adaptation patterns. Under monotherapy, tamox-ifen (A) predominantly induced transitions from ES to ER states (ES → ER), consistent with selection pressure favoring targeted therapy resistant clones. A small fraction of cells also transitioned toward mesenchymal-resistant (MR) phenotypes, suggesting limited EMT and immune escape under targeted molecular therapy (Fig 5C-D), which corresponded with notable tumor expansion (Fig 5K). ATRA (B) preserved epithelial states with minimal emergence of mesenchymal states (Fig 5C, E). In contrast, PD-1/PD-L1 blockade (C) exerted weaker selective pressure on ES states relative to tamoxifen, resulting in a less directed phenotypic response (Fig 5C, F). Despite inducing high T cell infiltration, both ATRA and PD-1/PD-L1 blockade alone were associated with substantial tumor expansion, indicating that immune access alone was insufficient for effective tumor suppression, potentially due to immune evasion mechanisms or a lag in cytotoxic response (Fig 5K). Dual therapies exhibited more complex dynamics. The combination of tamoxifen and ATRA (A+B) significantly reduced both ES → ER and ES → MR transitions (Fig 5C, G), while also maintaining strong T cell infiltration and limiting tumor growth, reflecting cooperative effects in sensitizing tumors to immune attack. Tamoxifen combined with PD-1/PD-L1 blockade (A+C) retained high ES → ER transition, indicating effective clearance of sensitive cells, but also allowed expansion of mesenchymal clones that evaded immune surveillance (Fig 5C, H). Tumor expansion was lower than with PD-1/PD-L1 blockade alone, likely due to tamoxifen-mediated elimination, though T cell infiltration was not further enhanced (Fig 5K). ATRA plus PD-1/PD-L1 blockade (B+C) preserved epithelial traits while enabling moderate immune susceptibility, producing a mixed phenotypic profile (Fig 5C, I) and intermediate tumor control (Fig 5K). Strikingly, the triple combination (A+B+C) minimized mesenchymal-associated transitions (ES → MR and ER → MR) and funneled tumor cells primarily into the ER state (Fig 5C, J). This group exhibited the lowest tumor expansion and sustained high T cell infiltration (Fig 5K), reflecting synergistic effects in constraining evolution and enhancing immune accessibility.

Together, these results demonstrate that treatment combinations differentially modulate tumor growth, phenotypic plasticity, and immune accessibility. Tumor adaptation shifts in both direction and magnitude in response to therapy, with multi-agent strategies exerting coordinated pressure to restrict malignant diversification, shape favorable spatial and phenotypic tumor landscapes, and promote effective immune engagement and spatial containment.

## Methods

### Gene Regulatory Network Simulation

The emergent dynamics of this GRN were simulated using the Random Circuit Perturbation (RACIPE) algorithm (20). RACIPE converts the network topology into a set of coupled ordinary differential equations (ODEs) and performs numerical simulations. For each simulation, the kinetic parameters are randomly sampled from biologically plausible ranges, allowing for an unbiased exploration of the system’s pheno-typic landscape. We generated an ensemble of models using 100,000 distinct parameter sets. For each parameter set, the ODEs were solved from 100 different initial conditions to identify all possible stable steady states.

The steady-state gene expression levels generated by RACIPE, which are provided on a log2 scale, were znormalized for comparative analysis. The z-score for each gene *Z*_*i*_ in a given steady state was calculated using the following equation:

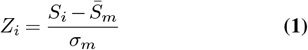

where *S*_*i*_ is the *log*_2_ expression value of the gene in that state, while 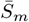 and *σ*_*m*_ are the mean and standard deviation of that gene’s expression across all calculated steady states from all 100,000 parameter sets, respectively.

To quantify the cellular phenotype of each steady state, we defined two metrics: an EM score and a Resistance score. The EM score was calculated as the normalized difference between mesenchymal and epithelial markers:

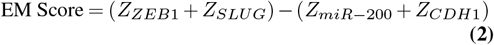

A higher EM score indicates a more mesenchymal phenotype. The Resistance score was defined based on the relative expression of the two ER*α* isoforms, with ER*α*36 being associated with resistance and ER*α*66 with sensitivity:

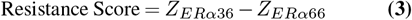

A higher Resistance score indicates a phenotype more likely to be resistant to anti-estrogen therapy.

### Analysis of Phenotypic Distributions

The parameter sets were first categorized based on the number of stable states they yielded (monostability, bistability, or tristability). For each category, the resulting steady states were classified into distinct phenotypes (e.g., Epithelial-Sensitive, Mesenchymal-Resistant, Hybrid E/M-Resistant) based on their positions in the two-dimensional space defined by the EM and Resistance scores. The distribution of these phenotypes was visualized using Kernel Density Estimate (KDE) plots. To investigate the link to immune evasion, the average z-normalized PD-L1 expression was calculated for each distinct phenotype within the monostable, bistable, and tristable cohorts. Error bars in the analysis represent the standard deviation of the corresponding z-normalized gene expression values.

### Agent-based model

We developed an off-lattice agent-based model based on the Gillespie algorithm to simulate tumor dynamics under varying environmental pressures, including adaptive T cell cytotoxicity and therapeutic intervention (24). The model comprises three main components.

a. Tumor behavior: We assumed a clonally homogeneous tumor population, in which all cells carried the same tumor antigens and no mutational heterogeneity was considered. Tumor cell events included division and migration. *Tumor division*. At each division step, parent cells were selected with equal probability. A daughter cell was then placed at a fixed distance from the parent in a randomly chosen direction. Division was accepted only if spatial constraints were satisfied, ensuring the daughter’s position did not overlap with the neighboring cells. Successful daughters inherited the phenotypic and kinetic attributes of their parent. *Tumor migration*. At each migration step, a live tumor cell was randomly chosen and proposed to move a fixed displacement according to a diffusion-based step size

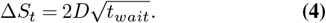
b. T cell behavior: All T cells were randomly divided into 500 clones, with each clone assigned a specific recognizable antigen. The tumor population was assumed to be homogeneous, consisting of a single tumor clone that expressed a repertoire of 100 antigens (24). Although in general sub-clonal expression of antigens can complicate tumor-immune interaction dynamics (24, 30, 31), we focused here on a clonal pattern of immune presentation so that each tumor cell was assumed to carry the same set of antigens, thereby allowing potential recognition by multiple T cell clones. All T cells were initially placed randomly along a circular boundary at a fixed distance from the tumor. *T cell migration*. At each migration step, a T cell was randomly selected to move. Without tumor cues, it performed a random diffusion-like step. When tumor cells were present, movement was biased toward the tumor centroid using a distance-weighted gradient (exponentially decaying with T-tumor distance with spatial decay constant *k* = 0.35). Candidate moves were accepted only if no overlap with other T cells occurred along the path, multiple intermediate waypoints were checked. *T cell killing and clonal expansion*. After a successful move, tumor cells within a fixed killing radius were identified. A target was randomly chosen from this neighborhood. If its antigen matched the moving T cell’s clone, the tumor cell was removed. The killing event also triggered T cell clonal expansion, adding a new T cell of the same clone at the target’s position.
c. Tamoxifen dynamics: Tamoxifen was modeled as a diffusing concentration field. The concentration evolved by diffusion and local uptake at tumor sites with:

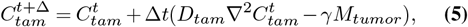

where *D*_*tam*_ is the diffusion coefficient, *γ* is the per-cell consumption rate, and *M*_*tumor*_ is the tumor occupancy mask. Tumor cells located at positions with *C*(*x, y, t*) ≥ *C*_*kill*_ were considered kill candidates. Their probability of death was given by

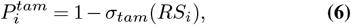

with *RS*_*i*_ the resistance score of cell *i*. For foundational understanding, *σ*_tam_ was taken as a sigmoid function represent-ing reduction in killing rate with increasing resistance score.

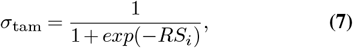
d. ATRA dynamics and phenotype conversions. To model ATRA effects within the ABM, we introduced a diffusing drug concentration field *C*_ATRA_(*x, y, t*). At each time step, the field was updated by finite-difference diffusion and local consumption at tumor-occupied grid sites:

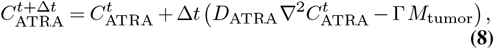

where *D*_ATRA_ is the diffusion coefficient, Γ the per-cell consumption rate, and *M*_tumor_ is the tumor mask matrix. For each tumor cell, we mapped its spatial location to the nearest grid index. Cells situated at positions with *C*(*x, y, t*) ≥ *C*_thresh_ were flagged as conversion candidates. Converted cells underwent EMT reprogramming according to GRN-informed phenotype assignments. Specifically, candidate cells were reassigned to new EMT states (E, H, or M) with corresponding updates to their motility rates, TamRes scores, EMT scores, and PD-L1 levels. These updates were drawn from precomputed GRN-derived phenotype-signature mappings.

### GRN-informed ABM initilization and phenotype transitions

Model parameters were chosen to ensure that the agent-based simulations produced tumor growth and phenotypic transition dynamics within biologically reasonable ranges, as supported by experimental observations and prior literature. In particular, the tumor division rate was sampled in the range [0, 2] with a uniform step size of 0.1, the tumor migration rate in the range [0, 1] with a step size of 0.05, and the T cell migration rate in the range [1, 2] with a step size of 0.2. These ranges were informed based on sensitivity analyses from prior work (24). Parameter sets that satisfied these criteria were subsequently incorporated into the GRN-driven simulations for further analyses. We then simulated the GRN to obtain an ensemble of steady states. From this ensemble, we randomly sampled *n* (*n* = 200) distinct states to seed the ABM. For each sampled state *s*, precomputed mapping dunctions were used to derive signature scores EMT, TamRes, and PD-L1. Cells were partitioned into E/H/M phenotypes based on the GRN-mapped scores. EMT phenotype-specific motility rates were assigned as

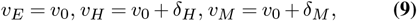

where *v*_0_ = 0.01 is the baseline motility rate, and *δ*_*H*_ = 0.3 and *δ*_*M*_ = 0.6 represent incremental motility offsets for H and M states, respectively. These parameters were selected to reflect the progressive increase in motility along the EMT axis, with hybrid cells displaying intermediate motility and mesenchymal cells exhibiting the highest motility (32). In this scheme, increasing EMT progression is reflected by correspondingly higher motility, with mesenchymal cells moving fastest and epithelial cells slowest (33, 34). To model progressive antigen loss along the EMT axis, we reduced the antigen repertoires for H and M from the E repertoire by random sub-sampling (34). EMT and TamRes scores were discretized to phenotype labels, yielding a combined label EMT × TamRes. These values initialized each cell’s three signature scores and its phenotypes in the ABM. Tamoxifen-induced death probability for a cell *i* was defined in Equation 7. Likewise, PD-L1-mediated immune evasion modulated T cell cytotoxicity via

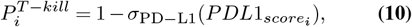

with 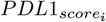 the resistance score of cell *i*. For foundational understanding, *σ*_PD*−*L1_ was taken as a sigmoid function representing reduction in killing rate with increasing PD-L1 score, with the coefficient applied as a scaling factor to enhance the sensitivity of the function to GRN-derived scores.

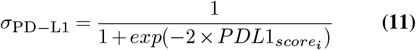

In non-plastic scenarios, phenotypes were fixed (no transitions). In plastic systems, we coarse-grained GRN outcomes into a finite set of phenotypes and modeled phenotype switching as a continuous-time Markov chain with generator *Q* = (*q*_*ij*_). Let *π* denote the stationary distribution estimated from GRN outputs (frequency of each phenotype across steady states). We constructed *Q* to satisfy *πQ* = 0 under detailed balance.

For the bistable (two-state) case (phenotypes *A* and *B*), the stationary distributions *π*_*A*_ and *π*_*B*_ were directly used from the GRN outputs, corresponding to the frequency of each phenotype across steady states. We fixed *q*_*AB*_ = *k*, with *k* = 0.5, and set 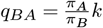 to satisfy detailed balance. The constant k was chosen as a biologically reasonable transition rate to ensure stable switching dynamics while preserving the GRN-derived equilibrium proportions.

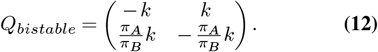

In the tristable scenario with phenotypes E, H, and M (three-state chain (E ↔ H ↔ M, without direct E ↔ M transitions)), the stationary distributions *π*_*E*_, *π*_*H*_, and *π*_*M*_ were used from the GRN outputs, representing the relative frequency of each phenotype across steady states. We fixed *q*_*EH*_ = *k* and *q*_*HM*_ = *ℓ*, with *k* = *ℓ* = 0.5, and the remaining rates were determined under the constraint of detailed balance. The constants *k* and *ℓ* were chosen as biologically reasonable transition rates to allow stable switching dynamics while preserving the equilibrium proportions dictated by the GRN-derived stationary distributions.

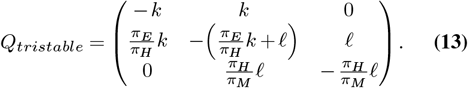

### Transcriptomic data processing and signature stratification

All the following computational analyses were conducted in the R environment. We queried the TCGA BRCA cohort from UCSC Xena. Only primary tumor samples were kept. Genes with *<* 1 expression in *>* 50% of samples or all missing values were excluded. ER+ samples were selected based on the ER status in clinical annotations.

To quantify phenotypic programs, we applied single-sample gene set enrichment analysis (ssGSEA). EMT scores were computed using the Hallmark EMT gene set from MSigDB (21). Tamoxzifen resistence (TamRes) scores were derived from a curated signature of upregulated genes identified in proteomic studies of resitant MCF7 cells, restricted to genes overlapping with the expression matrix. PD-L1 signatures scores were computed from a curated list of immune-related genes previously reported to correlate with PD-L1 activity (10). All scores were computed using the GSVA package with ssGSEA parameters. Samples were subsequently stratified into groups. EMT scores were divided into epithelial (E), hybrid (H), and mesenchymal (M) states using tertile cutoffs. TamRes and PD-L1 signature scores were both classified at the cohort median into sensitive (S)/resistant (R) and low/high groups, respectively. These classifications were further combined into a composite EMT × TamRes phenotype with six possible states (ES, ER, HS, HR, MS, MR). To evaluate relationships among the three phenotypic signatures, we performed pairwise correlation analysis. EMT, TamRes, and PD-L1 signature scores were extracted for all available samples. A pairwise scatterplot was generated with ggpairs.

ATRA-related gene signatures were derived from prior studies on transcriptomic changes in tamoxifen-sensitive (MCF-7) and -resistant (BT474) breast cancer cells treated with all-trans retinoic acid, curated into four sets: Responsive_*up*_, Responsive_*down*_, Resistant_*up*_, Resistant_*down*_. Next, ssGSEA was applied to the four sets to obtain per-sample enrichment scores. We then computed

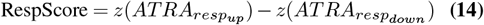

and

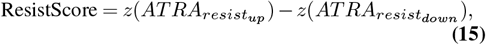

and defined

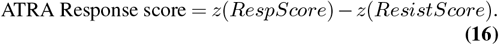

Samples were classified as Responsive-like or Resistant-like based on ATRA Response Score (*>* 0 vs *≤* 0).

## Discussion

Breast cancer remains one of the most prevalent malignancies worldwide, with its progression and treatment resistance shaped by both genetic and non-genetic mechanisms (35– 37). Among its subtypes, ER+ tumors pose a particularly unresolved challenge, as reversible resistance and immune evasion continue to undermine long-term therapeutic success (3). To understand the role of co-occurrence of reversible resistance and immune evasion on long-term therapeutic outcomes, we developed a multi-scale framework that connects a compact ER-EMT-PD-L1 regulatory circuit with a spatial tumor-immune ABM to examine treatment responses in ER+ breast cancer. Within this framework, we found that the ER-EMT-PD-L1 network generates multistable states that mechanistically link EMT to tamoxifen resistance and PDL1-driven immune evasion. Analyses of patient data reinforced the clinical relevance of these findings, demonstrating that EMT and resistance signatures coincide with elevated checkpoint expression and worse prognosis. Extending these correlations, examination of the ATRA-response signature indicated that patients with ATRA resistant-like profiles experience poorer survival, supporting the broader hypothesis that diminished MET-inducing capacity contributes to aggressive disease progression. Consistent with these clinical and molecular observations, ABM simulations further confirmed that tumor escape depends on the cooperative action of resistance, motility, and immune suppression, with plasticity and multistability accelerating these dynamics. Taken together, our results provide a rationale for integrated therapeutic strategies that combine endocrine therapy, MET inducers such as ATRA, and checkpoint inhibitors to restrict diversification and preserve immune access.

Our analysis revealed that the ER-EMT-PD-L1 regulatory circuit can sustain mono-, bi-, and tri-stable ensembles encompassing ES, hybrid E/M-resistant, and MR states, with PD-L1 levels rising stepwise across this spectrum. The coexistence of multiple stable states indicates that tumor populations can flexibly redistribute among epithelial, hybrid, and mesenchymal compartments in response to therapeutic or immune pressures, thereby accelerating relapse and resistance (38–40). Importantly, this stepwise trajectory underscores that immune evasion does not require a complete EMT; even partial EMT is sufficient to elevate PD-L1 expression and confer immune-evasive capacity (10, 41–43). In this way, our findings refine the conventional view that immune escape is largely confined to fully mesenchymal, invasive phenotypes, instead positioning the hybrid E/M state as a critical intermediate that couples proliferative potential with adaptive advantages (10). This interpretation is supported by prior theoretical and experimental work showing that hybrid states can exhibit immunosuppressive properties comparable to mesenchymal cells, mediated through ZEB1-miR200-PD-L1 circuitry and related regulatory axes (42, 44, 45). From a therapeutic perspective, the stepwise rise in PD-L1 expression suggests that endocrine resistance and immune evasion are not separate phenomena but co-emergent features of EMT plasticity. Consistent with this model, ssGSEA analysis across TCGA ER+ tumors demonstrated positive correlations among EMT, tamoxifen-resistance, and PD-L1 signatures, with hybrid and MR groups showing clear enrichment of PD-L1. These observations provide empirical validation of the modeled relationship and underscore important clinical implications: targeting estrogen signaling alone or PD-1/PD-L1 pathways alone may be insufficient, since hybrid states simultaneously link both mechanisms. Therapeutic strategies that either destabilize hybrid phenotypes or combine MET-inducing agents with checkpoint blockade may therefore represent more effective approaches to prevent tumor escape. More broadly, our results highlight PD-L1 as a continuous, rather than binary, readout of EMT progression, pointing to its potential utility as an early biomarker of adaptive immune evasion in ER+ breast cancer.

Building on these molecular insights, our ABM of tumor-immune interactions revealed that endocrine-resistant, EMT-positive, and PD-L1-high subpopulations adopt distinct yet complementary strategies of persistence. Resistant clones withstand therapeutic pressure, EMT-positive clones provide motility and invasive capacity, and PD-L1-high clones establish an immunosuppressive niche that protects the population. Rather than acting in isolation, these subpopulations cooperate as a functional collective, creating a division of labor that stabilizes tumor survival under various selective forces. Introducing phenotypic plasticity further amplified these dynamics, as cells could dynamically switch between states to restore weakened subpopulations, thereby accelerating escape and ensuring population adaptability. Thus, tumor progression emerges not from the dominance of a single pathway but from the synergistic interplay of resistance, motility, and immune suppression, reinforced by the buffering effects of plasticity and multistability.

This systems-level view highlights the limitations of approaches that target only one axis, such as endocrine signaling or immune checkpoint pathways in isolation. Effective intervention may instead require integrative strategies that simultaneously constrain resistance mechanisms, attenuate EMT-driven motility, and relieve checkpoint-mediated immune evasion, while also reducing the capacity for phenotypic interconversion that underlies rapid adaptation. In particular, our simulations indicated that a combined regimen of tamoxifen, cytotoxic T cell activity, and the MET inducer ATRA not only reduced mesenchymal-associated transitions (such as ES → MR and ER → MR) but also funneled trajectories toward the ER state. This three-pronged strategy produced the lowest tumor expansion and sustained T cell infiltration, suggesting a cooperative effect that constrained malignant diversification. These findings illustrate how targeting resistance, motility, and immune evasion in parallel can yield more durable control than single or dual therapies, and they highlight the potential of rationally designed combination regimens to improve immune accessibility. Comparable principles have been observed in breast and lung cancer where EMT, resistance programs, and immune evasion cooccur to sustain progression (46–49). These parallels highlight that tumor escape reflects not only the traits acquired but also the way those traits interact across space and time (27, 50–52). For instance, EMT-induced stromal remodeling not only drives motility but also alters immune infiltration patterns, while PD-L1 upregulation stabilizes these niches (53). Clinical pathology further shows that immune-excluded tumors often display stromal signatures and poor checkpoint response (54–57). Together, these observations imply that designing therapies requires more than combining drugs, it requires sequencing and spatial targeting to disrupt trait synergy and restore immune visibility. More broadly, these findings support an eco-evolutionary framework of tumor escape, in which clonal diversity, functional complementarity, and phenotypic plasticity collectively enhance resilience. In this light, endocrine resistance must be understood not only as a receptor-signaling defect but as an emergent property of a heterogeneous, adaptive disease, where spatial and temporal contexts further shape whether immune attack succeeds or is excluded.

Our framework emphasizes essential interactions, while some biological intricacies remain beyond its current scope. The GRN focuses on a minimal ER-EMT-PD-L1 module, while additional regulators, or metabolic and stress-response pathways, are not explicitly represented and could reshape attractor topology. The ABM similarly assumes clonal antigen uniformity and omits long-term genetic evolution, myeloid subsets, CAFs, Treg compartments, and T cell exhaustion, all of which are highly relevant to immune exclusion and checkpoint response. Further development of our framework will benefit from several important directions of investigations. Quantitative and spatial validation could combine single-cell RNA-seq with spatial transcriptomics or imaging mass cytometry in ER+ tumors to map EMT-PD-L1 gradients and T cell localization, thereby testing whether hybrid/MR niches coincide with immune exclusion. In parallel, targeted network perturbations using ZEB1/miR-200, ER*α*/ER*α*36, and LOXL2 in ER+ lines could reveal causal shifts in PD-L1 expression, MHC-I presentation, and susceptibility to T cell killing in autologous co-cultures, thereby testing the link between EMT programs, reduced antigen visibility, and immune evasion. Therapy sequencing also requires direct testing both *in vitro* and *in vivo* with time-staggered regimens, monitoring phenotypic shifts (EMT scores, PD-L1, apoptosis) and spatial T cell access. Moreover, ECM and biophysical interventions such as LOXL2 inhibition or collagen crosslinking blockade should be evaluated for their ability to relieve exclusion, particularly when combined with checkpoint inhibitors and ATRA. Finally, model extensions could incorporate macrophage and Treg modules, explicit exhaustion states, and mutational drift, calibrated to multiplex pathology, to capture eco-evolutionary interactions under chronic therapy.

Taken together, our findings highlight that endocrine resistance in ER+ breast cancer cannot be understood as a single signaling defect but as the emergent outcome of EMT plasticity, immune evasion, and their eco-evolutionary interplay. By linking molecular circuits to population dynamics and therapeutic responses, we show that durable tumor control will likely require multi-axis, context-aware strategies rather than monotherapies. This perspective underscores both the value of integrative modeling frameworks and the need for systematic validation to translate these insights into clinically actionable regimens.

## Declarations

### Availability of data and materials

The GRN-associated code is available at: https://github.com/csbBSSE/TamRes. The ABM-associated codes are available at: https://github.com/TAMUGeorgeGroup/EVO-ACT.git.

### Authors’ contributions

JTG and MKJ conceived of the research. YF and SS designed and performed the research. YF and SS developed the code. YF, SS, MKJ and JTG analyzed and interpreted the data. YF, SS, MKJ and JTG wrote the paper. All authors read and approved the final manuscript.

## Acknowledgments

We thank the Texas A&M High Performance Research Computing facility for providing advanced computing resources that supported portions of this work.

